# Sustainable *in situ* extraction of microalgae-derived terpenoids using functionalized silica microparticles

**DOI:** 10.1101/2024.08.26.609696

**Authors:** Sebastian Overmans, Adair Gallo, Himanshu Mishra, Kyle J. Lauersen

## Abstract

*Chlamydomonas reinhardtii* is a green microalga that has been genetically engineered to produce heterologous terpenoid metabolites. While this green biochemical approach for producing high-value chemicals holds tremendous potential, the current *in situ* extraction methods, utilizing traditional solvents such as dodecane and perfluorinated chemicals, present separation challenges, unfavorable economics, and risk of cell toxicity. Here, we develop a low-cost solvent-free approach based on silanized silica particles to fill the technology gap. We determine the feasibility and specificity of three differently coated (C11, C16, C18) silica particles to extract nine terpenoid metabolites from *C. reinhardtii* cultures during cultivation. The results reveal that the extraction efficiencies of functionalized particles, particularly those coated with C18, demonstrated high potential as an alternative to traditional extraction methods. While the extraction efficiencies of coated microparticles are for most compounds lower than those observed for dodecane-based extractions, it holds higher potential for larger-scale cultivations where the use of dodecane is restricted due to its flammability and tendency to form emulsions. In addition, the present study provides insights into the optimization of particle-to-culture volume ratio, particle saturation, and the required number of product elution steps, and proves the feasibility of an upscaled extraction of the sesquiterpenoid patchoulol as a representative metabolite in 5 L hanging bag reactors. This study paves the way for a circular bioprocess where product capture with traditional solvent overlays is impractical due to the formation of emulsions, and at the same time allows for an easy product recovery (solid-liquid separation) and reuse of functionalized particles over numerous cycles.

## 1 Introduction

Metabolic engineering of microbes and algae has emerged as a promising approach for the production of a variety of high-value terpenoid metabolites [1]. Metabolically engineered *Chlamydomonas reinhardtii* has been a significant player in this field due to its robust photosynthetic capabilities and the wealth of molecular tools available for its genetic manipulation [2, 3]. The green microalga’s well-characterized nuclear, chloroplastic, and mitochondrial genomes make it an attractive candidate for genetic engineering aimed at producing high-value bioproducts [4].

The production of high-value terpenoid metabolites using genetically engineered *C. reinhardtii* involves the introduction and expression of foreign genes responsible for terpenoid biosynthesis. For instance, a sesquiterpene synthase in *C. reinhardtii* was successfully expressed, leading to the production of valuable sesquiterpenes [5]. More recently, a novel approach was presented to recreate and concentrate complex mixtures of a wide variety of terpenoids typically found in fragrant agarwood [6]. These efforts highlight the potential of *C. reinhardtii* as a platform for the sustainable production of bioactive compounds. However, a persistent challenge remains in efficiently extracting these metabolites *in situ*.

Traditionally, dodecane overlays have been employed to extract metabolites directly from the culture medium due to the solvent’s ability to sequester hydrophobic compounds effectively [3, 7, 8]. However, dodecane presents several limitations, including difficulties in separation from the culture medium, the formation of micelles, and potential toxicity to the algal cells at higher concentrations [9, 10]. Recent studies have explored the use of fluorinated compounds (FCs) for in situ extraction of sesquiterpenoid metabolites from algal cultures [6, 11]. FCs offer advantages over dodecane overlays, such as ease of separation from the culture and compatibility with liquid-liquid extraction processes using solvents like ethanol [12]. Despite these benefits, FCs are associated with issues such as evaporation and environmental persistence [13], although new methods for their degradation have recently been proposed [14, 15].

To address these challenges, we considered the application of surface engineered solid hydrophobic particles for extracting metabolites from algal cultures, which could be separated easily via sedimentation and/or sieving. We hypothesized that if the chemical make-up of the particles is rendered similar to the water-dodecane interface, the consequent intermolecular and surface forces may afford metabolite physisorption on the particles [16–20]. Previously, water–hydrophobe interfaces have been exploited for several practical applications, such as non-stick surfaces [21], concentrating fruit juices via vacuum membrane distillation [22], treating wastewater [23, 24], suppressing water evaporation from irrigated soils (mulching) [25, 26], long-term seed storage [27], among others. In this study, we explore the feasibility of using low-cost silica microparticles with three different functional coatings to extract heterologous terpenoid metabolites from cultures of metabolically engineered *C. reinhardtii* in situ. We utilized nine strains of *C. reinhardtii*, each producing a different terpenoid metabolite, and compared the extraction efficiency of these functionalized microparticles against traditional extraction methods using dodecane and FC-3283. Initially, key parameters such as particle saturation and the required particle-to-culture ratio were investigated before we assessed the potential to upscale this cultivation- and extraction method using 5 L hanging bag reactors. Through this approach, we aim to provide a more sustainable and efficient method for in situ extraction of valuable metabolites from microalgal cultures.

## 2 Materials and Methods

### 2.1 Chemicals

The terpenoid extraction potential of functionalized microparticles was compared against the extraction efficiency of other chemicals frequently used for microalgal two-phase cultivations. Specifically, the two compounds selected as extraction baseline were the commonly used straight-chain alkane *n*-dodecane (≥99%, VWR International, France), which forms an overlay above algal cultures and is known to continuously “milk” terpenoid metabolites from living cultures [28]. Before being added to the algal culture, *n*-dodecane was filtered using a Supelclean^TM^ LC-Si solid-phase extraction column (Product no. 505374; Sigma-Aldrich, Germany) to remove impurities. The second compound against which we compared the extraction potential of microparticles was the perfluorinated amine FC-3283 (Acros Organics, Belgium), which also acts as a physical product sink but forms a dense underlay when added to microalgal cultures [12]. After cultivation, the particles were washed with 96% vol. ethanol (VWR International, France) to elute any adsorbed terpenoid from the functionalized microparticles.

The silanes used in the study were C18 – octadecyltrichlorosilane (Sigma-Aldrich, Germany), C16 – hexadecyltrimethoxysilane (Sigma-Aldrich, Germany), C11 – 10-Undecenyltrichlorosilane (Gelest, USA). Additionally, the following chemicals were used in the silanization protocol: acetone (Thermo Fisher Scientific, UK), sulfuric acid (VWR International, USA), hydrogen peroxide (Sigma-Aldrich, Germany), toluene (VWR International, USA), hexane (Sigma-Aldrich, Germany).

### 2.2 Description, functionalization, and E-SEM visualization of microparticles

The particles used in the present study were hydrophilic, silica-based Supelclean LC-SI microparticles with a nominal particle size of 45 µm and a surface area of 475m^2^/g (Product no. 57200; Supelco^TM^, Bellefonte, PA, USA). Some of those particles were functionalized with straight alkane chain silane groups to render them hydrophobic (Fig. 1a) following a previously described protocol [29]. Briefly, the particles were washed with deionized water to remove surface contaminants, then stirred in a beaker with acetone, filtered and rinsed with water and dried at 100°C for >6 h in an oven. Subsequently, the particles’ surface was activated in a freshly made piranha solution (3:1 volume ratio of 99.9% sulfuric acid and 30% hydrogen peroxide) for 10 min at 130°C. The activation step created hydroxyl groups on the surface of particles, which served as the binding sites for the silane molecules. Then, the piranha solution was drained followed by thoroughly rinsing the particles several times with deionized water. Next, the particles were dried at 120°C for ∼2.5 hours and cooled down. Immediately after, silanization reactions were performed with ratios of 5 g of particles to 2mL of the required silane in 8 mL of toluene in a stirred beaker at 40°C and for 3 h. The particles were subsequently rinsed in toluene several times followed by hexane to remove the unreacted silanes, then oven dried at 100°C, and stored in glass vials. Environmental scanning electron microscopy (E-SEM, Quattro, Thermo-Fisher) was used to qualitatively demonstrate the enhancement of silica particles’ surface hydrophobicity after silanization (e.g., C18). Sample preparation entailed sticking them to the stub via a double-sided copper tape. After the samples were introduced in the E-SEM chamber, its temperature was lowered via a Peltier stage and maintained at 293 K. Next, moisture was introduced in the chamber by raising its relative humidity, leading to ≈750–800 Pa. Images were captured using a secondary electron detector at 30 kV accelerating voltage, beam current 39 pA and a working distance of 4–5 mm.

**Figure 1.**
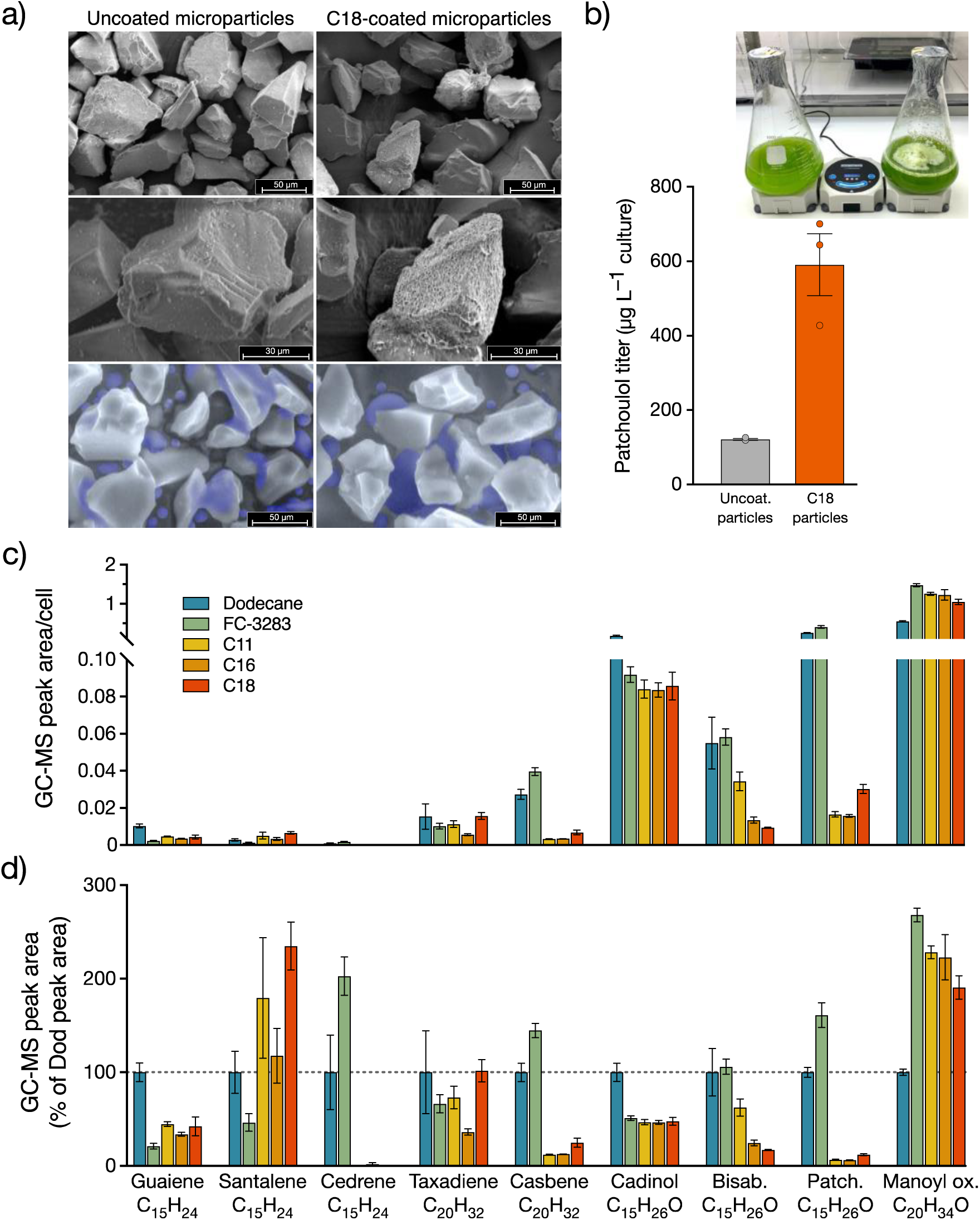
a) E-SEM visualization of uncoated-(left panels) and C18-coated (right panels) Supelclean microparticles. In the two lower panels, condensed water is artificially colored in purple. **b)** Patchoulol titer after 7-day cultivation of patchoulol-producing *C. reinhardtii* strain with either uncoated- (left flask) or C18-coated microparticles (right flask). Extraction efficiency of assorted algae-produced terpenoids when cultivated with dodecane, FC-3283, or differently coated microparticles, with extraction **c)** expressed as normalized by cell or **d)** compared against dodecane extraction efficiency. Error bars indicate standard error (SEM) based on three replicate measurements (n=3).

### 2.3 Microalgal strains

The algae that were used in the present study to determine the terpenoid extraction potential of functionalized microparticles when added to living terpenoid-producing microbial cultures were all genetically modified strains of the green microalga species *Chlamydomonas reinhardtii*. A total of nine *C. reinhardtii* strains were selected that had previously been engineered to either produce one of the diterpenoids taxadiene, casbene, and 13R(+) manoyl oxide [30], or one of the sesquiterpenoids patchoulol [31], ∂-guaiene, santalene, cedrene, cadinol, or bisabolol [6].

The algal strains were routinely maintained on agar plates consisting of Tris-acetate phosphite medium with nitrate (TAPhi NO3) [31], except the three strains producing taxadiene, casbene, and 13R(+) manoyl oxide, which were grown on agar plates made of the phosphate-containing variety of Tris-acetate medium with ammonium (TAP NH4) [32]. All cultures were grown under 50 µE PAR intensity before being transferred into 24-well microtiter plate wells containing 1mL TAP medium, and left shaking at 190rpm under 12 h: 12 h light: dark (150 µE PAR) illumination at 25°C. After 48 h, 500 µL of medium was added to each well to replenish nutrients. By day 4, 400 µL of each culture was inoculated in 40 mL growth medium in Erlenmeyer flasks and remained at 120 rpm agitation under the light conditions mentioned above until being used for subsequent experiments.

### 2.4 Determining compound extraction specificity of differently functionalized **microparticles**

To identify the compound extraction specificity of different particle coatings, we grew nine different *C. reinhardtii* strains (see description in 2.3) with either dodecane, FC-3283, or Supelco microparticles coated with C11, C16, or C18 chains.

Specifically, 200 µL of each strain was transferred from the 400 mL pre-cultures into individual microtiter wells of 6-well plates containing 4.3 mL growth medium. To each well, either 500 µL of dodecane or FC-3283, or 250 µL of microparticles were added, and the cultures were grown at 25°C and 120rpm agitation under 12 h: 12 h light: dark (150 µE PAR) illumination. The patchoulol-producing strain was additionally grown in 1 L Erlenmeyer flasks with a culture working volume of 400 mL together with 20 mL of either uncoated- and C18 Supelco microparticles under the above conditions, except agitation was provided by a stir bar (120 rpm).

After 7 days, 10 µL of culture was sampled from each replicate microtiter plate well and used to enumerate cell concentration using flow cytometry as outlined below. The remaining content of each well was transferred into individual 15 mL Falcon tubes. Samples with dodecane and FC-3283 were centrifuged at 4000 x *g* for 5 mins, and the dodecane overlays and FC-3283 underlays were recovered and pipetted into individual clean 2 mL Eppendorf tubes, centrifuged at 12,000 g for 5 mins, and the patchoulol content in the solvent quantified using GC-MS (see section 2.10). For cultures with microparticles, the tubes were left for 15 mins to allow the particles to settle. The supernatant consisting of culture and media was discarded, and the microparticles were washed with 2 mL of 96% ethanol, left shaking at 190 rpm for 2 h, centrifuged at 4000 x *g* for 5 mins, and the content of patchoulol in the ethanol quantified using GC-MS. Each co-cultivation condition was set up in biological triplicate wells (n=3).

### 2.5 Optimizing particle-to-culture volume ratio

To determine the lowest volume of C18-coated microparticles required to get the best possible extraction of the secondary metabolite patchoulol, we grew the patchoulol-producing *C. reinhardtii* microalgal strain with three different amounts of particles.

In per-autoclaved 500 mL bottles, three different volumes (237.5 mL, 243.75 mL, 247.5 mL) of low cell density *C. reinhardtii* culture (∼1×10^5^ cells mL^−1^) were grown together with either 12.5 mL (5%), 6.25 mL (2.5%) or 2.5 mL (1% of total volume) of C18-coated microparticles to achieve a total volume of 250 mL (culture + particles) in each bottle. Each condition was set up in five biological replicates (n=5). The cultures were grown for 7 days at 25°C and 12 h: 12 h light: dark (150 µE PAR) illumination under constant sparging with 3% CO2 in air (72 mL min^−1^) to keep the particles in suspension (see experimental setup in Fig. 2B). After 7 days, the sparging was stopped and left for 20 mins to allow the particles to settle. The supernatant consisting of culture and media was discarded. The microparticles of each bottle were transferred into a separate clean 50mL Falcon tube, washed with 25 mL of 96% ethanol, and shaken at 80 rpm for 15 mins before the entire ethanol was recovered for further analysis. This wash step was repeated two more times, totaling three wash steps per replicate culture. The content of patchoulol in the ethanol wash fractions was quantified using GC-MS (see section 2.10).

**Figure 2.**
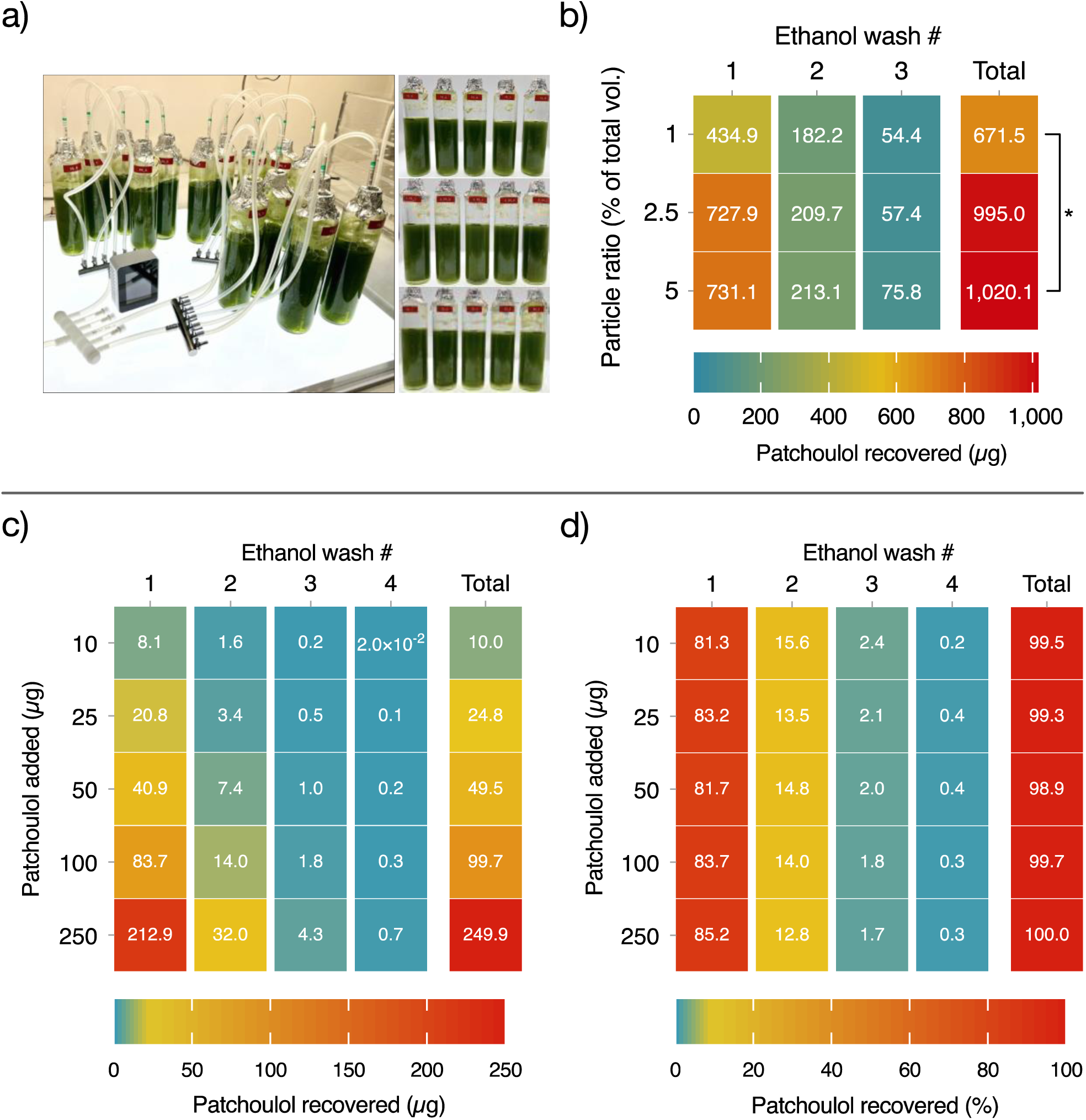
Results of the particle ratio experiment (a, b) and the experiment to determine the required number of product elution steps (c, d). **a)** Photographs of the experimental setup on day 6 (left panels) and the five replicate culture bottles per treatment (n=5) after 7 days of cultivation (right panel) **b)** Mean patchoulol extracted from *C. reinhardtii* culture during 7-day cultivation with C18 functionalized microparticles added into the culture at a ratio of either 1%, 2.5% or 5% of the total culture volume. The asterisk indicates the instance where the Student’s *t*-test identified significantly different mean values (*p* <0.05). Recovery of patchoulol standard from microparticles coated with C18 functionalized groups, expressed as **c)** absolute amount recovered (in µg) and **d)** percentage recovered based on initial amount added, shown for each of four wash steps of the particles with 96% ethanol. Values in c) and d) are means of three replicates (n=3).

### 2.6 Determining particle saturation and the required number of product elution **steps**

To identify the number of ethanol washes required to recover ≥99% of patchoulol product from C18-coated particles, known amounts of patchoulol standard were added to the particles and washed several times with ethanol.

Specifically, solid patchouli alcohol standard (Cayman Chemical Company, USA) was dissolved in 96% vol. ethanol and serially diluted to prepare five different concentrations (200, 500, 1000, 2000, 5000 ppm) of standard. 50 µL of each standard was added into individual Eppendorf tubes containing 950 µL of TAPhi NO3 medium. Each standard was prepared in triplicates (n=3). 50 µL of C18-coated microparticles (dry bulk density: 0.45 g/cm^3^) were added to each tube, then shaken at 800 rpm for 1 h, and centrifuged at 8000 x *g* for 5 mins before the supernatant was discarded.

The pelleted particles in each tube were washed with 500 µL of 96% ethanol, shaken at 800 rpm for 1 h, and centrifuged at 8000 x *g* for 5 mins. The supernatants were transferred into individual Eppendorf tubes and kept to quantify patchoulol concentration. This ethanol wash cycle of the particles was repeated three times to get a total of four washing cycles. All ethanol supernatants were centrifuged (8000 x *g*, 5 mins) and transferred into amber GC vials for quantification of patchoulol using GC-MS as described below. The amount of patchoulol in each sample was reported as both µg of patchoulol recovered and % recovered compared to patchoulol added.

### 2.7 Upscaling of patchoulol extraction in hanging bag reactors

To test the feasibility of using functionalized microparticles as product sinks in upscaled bioreactors, we grew the patchoulol-producing *C. reinhardtii* strain in commercially available 10 L hanging bag reactors. Specifically, three replicate hanging bags (n=3) were filled with 5 L of T2Phi medium and 100 mL of dense *C. reinhardtii* culture. 100 mL of C18-coated microparticles were added to each bag, and the reactors were kept at 25°C under constant illumination from five light panels (235 µE PAR) and sparging with 3% CO2 (300 mL/min) for 7 days (see setup in Fig.3 a).

**Figure 3.**
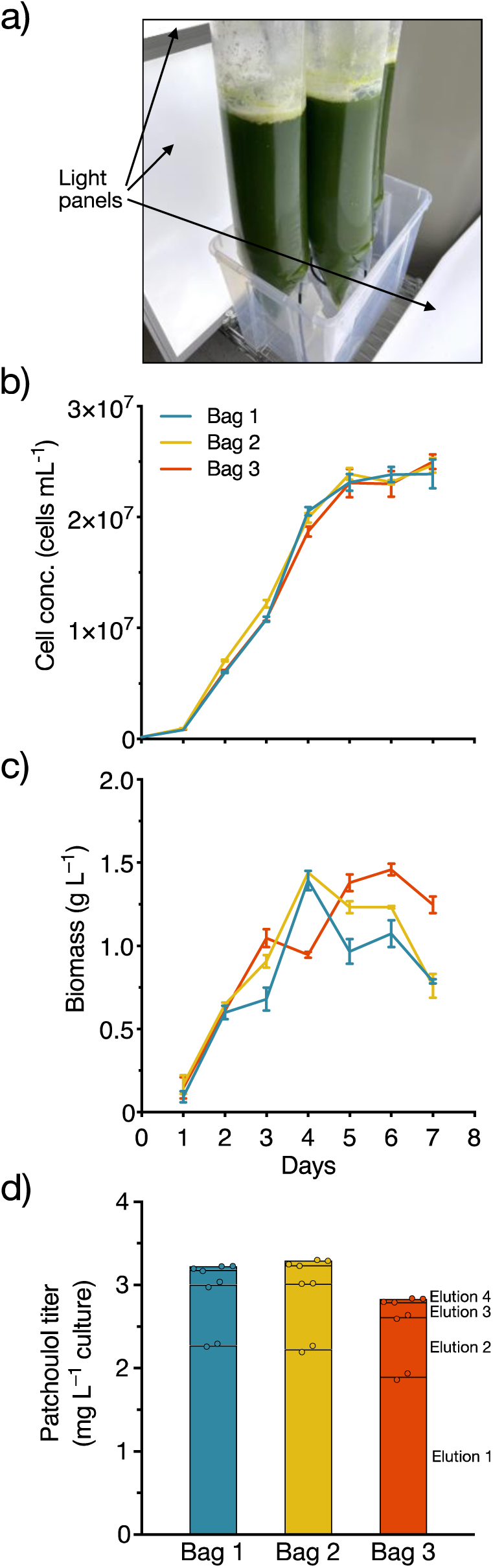
Upscaling of patchoulol production in hanging bags. a) photograph of the experimental setup on day 6, showing the three hanging bags, each containing 5 L of dense *C. reinhardtii* culture and 100 mL of C18-functionalized microparticles, and the light panels mounted on either side of the bags. b) *C. reinhardtii* cell concentration and c) algal biomass in three hanging bags over the 7-day experimental period; d) Patchoulol titer after 7 days of cultivation, shown for each bag and elution step. Elutions 1–4 refer to the four rounds of patchoulol product elution from the particles performed with 96% ethanol.

**Figure 4.**
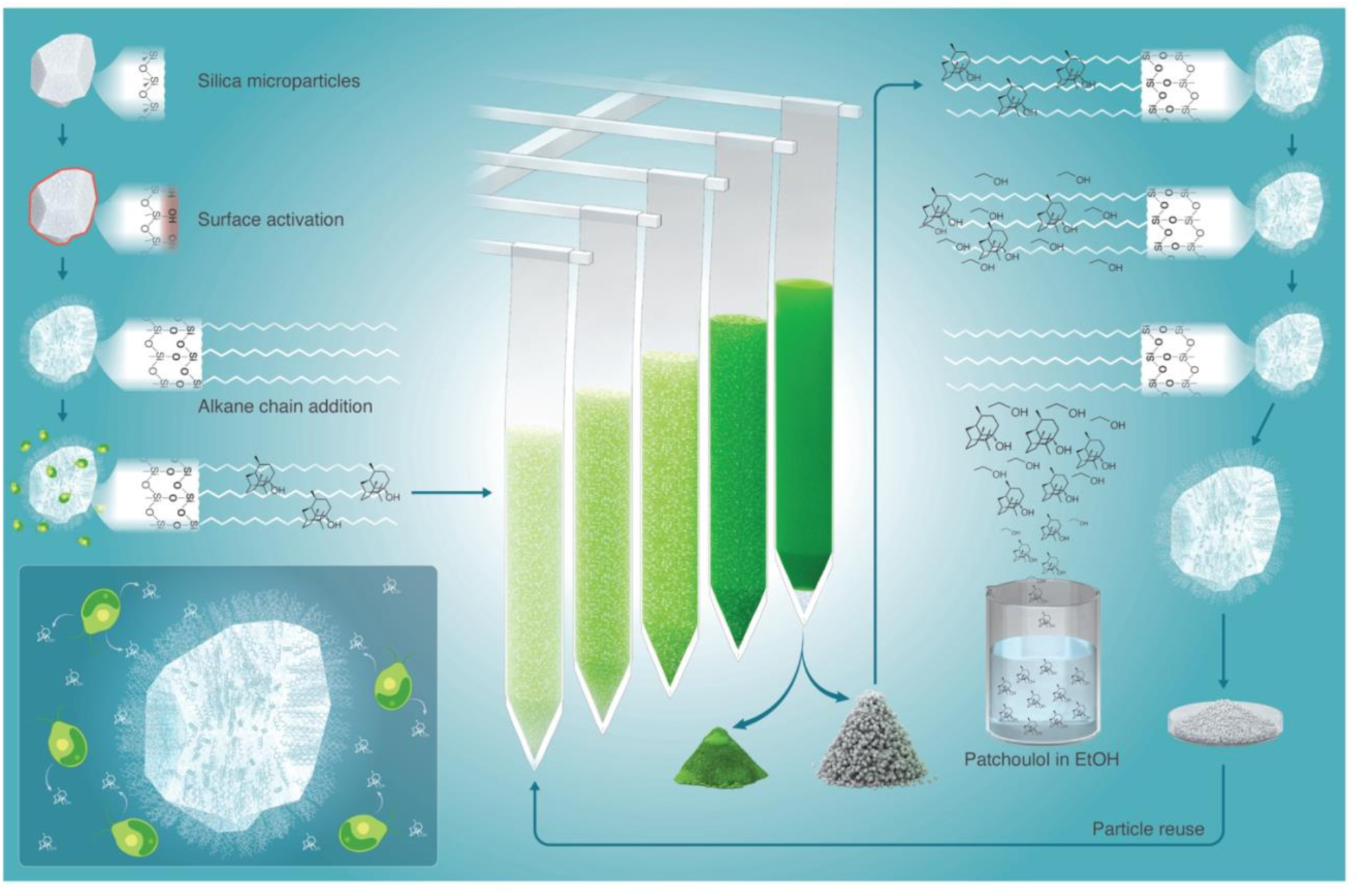
Illustration of the workflow used to functionalize microparticles and to extract the terpenoid product patchoulol from living microalgae cultures grown in hanging bag reactors. Left panels: Surface of silica-based microparticles gets activated before alkane chains are added to the surface in the process called silanization, and the coated particles are added to microalgal cultures. Middle panels: The coated microparticles get transferred to hanging bags containing a light green (low cell concentration) culture of a patchoulol-producing *C. reinhardtii* strain that gets continuously sparged with air and 3% CO2 to keep algal cells and particles in suspension. The algae are grown for 7 days until cultures are dark green in color due to high cell concentrations, at which point the algal biomass and particles can be recovered from the hanging bag and separated from each other. Right panels: The particles get washed with 96% ethanol to elute any patchoulol bound to the alkane chain coating. The end product is patchoulol diluted in ethanol, while the now clean, product-free particles can be dried and reused for further microalgal cultivations. Figure created by Heno Hwang, scientific illustrator.

20 mL of culture were sampled daily, left on the bench for 5 mins to allow the particles to sediment, and then used for flow cytometry and biomass determination (see details below).

After 7 days, the contents of each bag were transferred into individual 5 L measuring cylinders, and left for 15 mins until the microparticles had sedimented. The supernatant consisting of algal culture was discarded, while the microparticles were 4 x washed with 200 mL of ethanol as described above. 1 mL of each ethanol wash sample was centrifuged (8000 x *g*, 5 mins) and transferred into amber GC vials for patchoulol quantification using GC-MS.

### 2.8 Flow cytometry

Growth of algal cultures was monitored using a previously described protocol with slight modifications (Overmans and Lauersen, 2022). Cell densities were measured using an Invitrogen Attune^TM^ NxT Acoustic Focusing Cytometer (Thermo Fisher Scientific, UK). 15 µL of each biological replicate sample was diluted in 1485 µL of 0.9% NaCl solution (1:100 dilution) in technical triplicate (n=3) Eppendorf tubes. Immediately prior to the analysis, each sample was briefly vortexed. The first 25 µL of each sample was discarded to ensure a stable flow rate of cells during the measurement. Data acquisition stopped automatically when 50 µL from each tube was analyzed. All post-acquisition analyses and population gating based on forward scatter and chlorophyll autofluorescence were performed using the Attune^TM^ NxT Software v3.2.1 (Life Technologies, USA).

### 2.9 Dry biomass quantification

For the hanging bag experiment, *C. reinhardtii* biomass was determined daily by transferring 5 mL of culture into a pre-weighted test tube. The tubes were loosely covered with aluminum foil, left in a 110 °C oven for 48 h, and allowed to cool for 10 mins at room temperature before weighing the test tubes containing the dry biomass. Additionally, on the first day, 5 mL of fresh T2Phi medium was transferred to a pre-weighted test tube and processed in the same way as algal cultures. The weight of the dried medium was subtracted from the dry weight of each culture sample. All dry-weight analyses of cultures and media controls were performed in technical triplicates (n=3).

### 2.10 Gas chromatography-mass spectrometry (GC-MS) analysis

All samples were analyzed using an Agilent 7890A gas chromatograph (GC). The instrument was equipped with a DB-5MS column (Agilent J&W, USA) attached to a 5975C mass spectrometer (MS) with a triple-axis detector (Agilent Technologies, USA). A previously described GC oven temperature protocol was used [12]. After a 13 min solvent delay, mass spectra were recorded in scan mode in the range 50–550 *m/z* at 20 scans s^−1^. Chromatograms were processed and integrated using MassHunter Workstation software v. B.08.00 (Agilent Technologies, USA), and metabolites were identified by comparing their mass spectra against the National Institute of Standards and Technology (NIST) library (Gaithersburg, USA).

Chromatograms and peak area integrations were manually inspected for quality control. All GC-MS measurements were performed in technical duplicates (n=2).

### 2.11 Data analysis

Individual Student’s *t*-tests were performed to identify if mean values of total extracted patchoulol were significantly different (*p* < 0.05) between algal cultivations with C18 microparticle volumes of 1%, 2.5%, and 5%. JMP Pro 16.2.0 (SAS Institute Inc., Cary, NC) was used for data analyses, and GraphPad Prism v.10.2.3 (GraphPad Software, USA) for data visualization. Visual elements were harmonized in Affinity Publisher v.1.10.8 (Serif Ltd., West Bridgford, UK).

## 3 Results & Discussion

### 3.1 Metabolite extraction specificity of differently coated microparticles

The initial co-cultivation experiment with patchoulol-producing *C reinhardtii* grown together with either uncoated- or C18-coated microparticles in 1L Erlenmeyer flasks found considerable differences in extraction efficiencies. Specifically, uncoated particles extracted 121 ± 2.3 µg patchoulol/L culture (mean ± SEM), while C18-coated particles extracted almost five times more patchoulol (592 ± 70.2 µg/L culture) (Fig. 1b) due to the altered properties of the particles such as hydrophobicity (Fig 1a).

When we compared the terpenoid extraction potentials of dodecane, FC-3283, and different particle coatings, we found vast differences between solvents and coatings. As mentioned in previous studies, the algal strains used here are known to have a wide range of terpenoid product titers due to distinct metabolic engineering strategies and efforts (see Fig. 1c) [6, 30, 31], so we normalized the extraction against the amounts of product extracted by dodecane, which is frequently used for extractions of the sesquiterpenoid patchoulol from microbes [33, 34], including the microalga *Chlamydomonas reinhardtii* [3], and is known to have higher extraction efficiencies than perfluorinated compounds [12].

For the compounds produced by the nine *C. reinhardtii* strains used here, there was no clear extraction efficiency pattern based on the number of carbon atoms or the presence/absence of oxygen in the terpenoid product molecule. Guaiene and cadinol were most effectively extracted with dodecane (Fig. 1c), while the extraction with FC3283 or C18-coated particles was ∼50% lower. Cedrene, casbene, bisabolol, patchoulol, and manoyl oxide extraction titers were highest with FC3283. Santalene and taxadiene extraction was most efficient with C18-coated microparticles, whereas almost no cedrene extraction was achieved with either of the coated particles. When comparing particle coatings, the shortest chain coating (C11) was most efficient for the extraction of guaiene, bisabolol, and manoyl oxide, while the longest chain coating (C18) was best at accumulating santalene, taxadiene and patchoulol. For bisabolol and manoyl oxide, it was apparent that extraction efficiencies gradually decreased with increasing chain length of the particle coating. While the extraction efficiencies of coated particles were lower compared to dodecane for certain metabolites, microparticle-based product extraction would allow for upscaled microalgal cultivations where traditional alkane solvent overlays are impractical due to the formation of emulsions [10]. Additionally, this method shares the same benefits as *in situ* product extraction with FC-3283, such as biocompatibility, ease of product recovery using liquid-liquid extraction with ethanol, and that the extraction medium can be reused without the need for additional preparation steps [12]. Although, it is noteworthy that cell densities of cultures with microparticles are typically lower than those of cultures with dodecane overlays (Suppl. Fig 1), likely due to increased shading [35, 36], as well as shear stress during culture agitation [37] when microparticles come in contact with cells of the different *C. reinhardtii* strains that are all based on the cell-wall deficient UVM4 strain [38].

### 3.2 Elution- and particle volume-dependent extraction efficiency

Following the initial comparison of terpenoid extraction from different algal strains with various particle coatings, we used the highly engineered patchoulol-producing strain and assessed product extraction using different volumes of C18-coated particles. Subsequently, using a commercial patchoulol standard, we determined the number of ethanol washes required to recover 99% of the product from the particles.

The 7-day cultivation of the patchoulol-producing *C. reinhardtii* strain in sparged (3% CO2) bottles together with C18 microparticles at either 1%, 2.5%, or 5% of the total volume (Fig. 2a) revealed that across the different particle volumes, of the total extracted patchoulol, 65–73% was eluted during the first ethanol wash, 21–27% in the second wash and 6–8% during the third wash step (Fig. 2b). The mean amount of total extracted patchoulol gradually increased with particle volume (Fig. 2b), however, only the total amounts extracted with 1% particles (672 µg) and 5% particles (1020 µg) were significantly different (*t*(12)= 2.303, *p* < 0.05), while the difference in mean patchoulol recovered between 1% particles and 2.5% particles (995 µg) was not quite significant (*t*(12)=2.138, *p* = 0.0538). No difference in patchoulol extraction was found between 2.5% and 5% particles (*t*(12)=0.166, *p* = 0.0871). Given the fact that 5% particle volume did not result in significantly higher patchoulol extraction, and that lower cell densities were observed with 5% particle volume at the end of the experiment (Suppl. Fig. 2), likely due to the abovementioned increase in shading and shearing effects [36, 37], we decided to use a particle volume of 2% for subsequent cultivations in the hanging bag reactors.

To confirm that the particles’ surface area is sufficiently large to not get saturated with patchoulol during algal cultivations, and to determine how many ethanol elution steps are necessary to recover adsorbed patchoulol, known amounts of patchoulol standard were added medium suspensions with 50 µL of C18-coated microparticles. Overall, we identified that the particles were able to adsorb even the highest amount of patchoulol added (250 µg) (Fig. 2c), implying that particle saturation was not reached. Across the different treatments, we could observe that the bulk of patchoulol was eluted during the first ethanol wash (81–85%). During the second and third wash, 13–16% and 2% of the product were eluted, respectively, and only < 0.4% of the added patchoulol standard was recovered during a fourth wash (Fig. 2d). Irrespective of the added patchoulol amounts (10–250 µg), four ethanol elution washes were sufficient to elute ≥99% of patchoulol from the particles. As such, we opted to perform four product elutions with the particles recovered from the hanging bag reactors. To further concentrate the recovered terpenoid product, an energy-efficient organic-solvent nanofiltration concentration process could be used [6, 39], although no further product concentrations were performed here.

### 3.3 Upscaling of particle-based patchoulol extraction in hanging bag **bioreactors**

The patchoulol-producing *C. reinhardtii* strain was grown together with C18-coated microparticles in three illuminated hanging bag bioreactors that were continuously sparged with a 3% CO2 in air mix to provide a carbon source for the algae and keep the particles in suspension (Fig. 3a). During the seven days of cultivation, we observed that, in all replicate reactors, cell densities were increasing exponentially until day 4 and then reached a stationary phase, with maximum concentrations of 2.5×10^7^ ± 5.7×10^5^ cells mL^−1^ (mean ± SD) by day 7, which is lower than cell densities reported for this *C. reinhardtii* strain when grown in a high-cell-density CellDeg reactor with a dodecane overlay (∼1×10^8^ cells mL^−1^) [31]. However, the final cell densities here are slightly higher than the concentrations of 2.2×10^7^ ± 4.4×10^6^ cells mL^−1^ recorded for the sparged bottles cultivation setup with 2.5% particles (Fig. 2a; Suppl. Fig. 2). This finding and the consistent and stable culture growth observed across the three replicate bags indicates that engineered *C. reinhardtii* strains can be grown with coated microparticles in this reactor style. Algal biomass (g L^−1^), on the other hand, varied considerably between the three replicate reactors. For reactors 1 and 2, we observed the highest biomass was reached on day 4 (1.39 g L^−1^, 1.44 g L^−1^) and gradually decreased the following days, while for the third reactor, maximum biomass was measured on day 6 (1.46 g L^−1^) (Fig. 3c). Although it is noteworthy that this discrepancy between replicate reactors could stem from an incomplete separation of algal biomass and microparticles in certain samples, with minor traces of particles adding additional weight to the biomass measurements, whereas the flow cytometric analysis only records fluorescent algal particles.

The recovery of patchoulol product from the microparticles on day 7 of the experiment revealed a maximum patchoulol titer of 3.30 mg L^−1^ culture in reactor 2, and an average titer of 3.12 ± 0.25 mg L^−1^ culture across the three reactors (Fig. 3d), supporting the claim that this novel particle-based terpenoid metabolite extraction method can be upscaled in this reactor style and provide consistent product recovery. The average patchoulol titer here is over three times higher (0.84 mg L^− 1^) than that reported for a perfluorocarbon-based patchoulol extraction from *C. reinhardtii* cultures grown on AnMBR effluent in 400 mL Algem reactors for 4 days [11]. However, the titers here are only ∼50% compared to the titers reported for *C. reinhardtii* cultures grown in CellDeg HD100 cultivators with dodecane overlay (6.2 mg L^−1^)[31]. As such, future studies could explore the feasibility of using functionalized microparticles in high-density cultivators such as the CellDeg system.

### 3.4 Particle stability and reuse

In addition to the benefits noted above, we can enhance the process sustainability if the functionalized silica particles could be reused over numerous cycles. Silanized silica surfaces are known to degrade over time when they are subjected to elevated temperatures, harsh pH swings, or mechanical abrasion [40, 41]. Secondly, hydrophobic surfaces are known to be vulnerable to organic fouling in the presence of amphiphilic molecules, which leads to wetting reversal and loss of function [42, 43]. In either scenario, surface wettability drops monotonically over time and so does the process efficiency. We note that our process is devoid of harsh physical conditions, which guarantees long-term stability of the surface chemical make-up. Wetting reversal due to organic fouling of particles is not concern for our process because we utilize ethanol wash as the termination step of the process, wherein physisorbed terpenoids dissolve into the solvent, thus regenerating the functionalized particles. Notably, silane bonds at the particle surface are strong enough (bond strength ∼100 kcal/mol) to withstand the molecular forces during attachment/detachment of cycles of terpenoids (non-bonding interactions ∼2–10 kcal/mol), highlighting the high reusability potential of C18 terminated particles for numerous product accumulation and recovery cycles.

### 3.5 Conclusion

Our study aimed to determine the feasibility and specificity of differently functionalized microparticles to extract heterologous terpenoid metabolites from metabolically engineered *C. reinhardtii* strains *in situ* during cultivation. The results showed that functionalized microparticles, particularly those coated with C18, demonstrated potential as an alternative to traditional extraction methods, such as dodecane overlays and underlays of perfluorinated compounds such as FC-3283, for the extraction of terpenoid metabolites from living microalgal cultures. Additionally, the study provided insights into the optimization of particle-to-culture volume ratio, particle saturation, and the required number of product elution steps, as well as proved the feasibility of an upscaled extraction of patchoulol as a representative metabolite in hanging bag reactors. While the extraction efficiencies of coated particles were for some compounds lower compared to dodecane, the particle-based method could enable upscaled microalgal cultivations where product capture with a traditional solvent overlay is impractical due to the formation of emulsions. Notably, only a tiny fraction (< 0.1%) of hydrocarbon/perfluorocarbon chemicals in our approach, compared to the amount of chemicals exploited in the traditional approach. The ease of product recovery and particles’ reusability underscore the practicality and sustainability of this circular process. As regulations on the use of PFCs in processes rise globally [44], this low-cost, simple, and robust process should present a sustainable alternative.

## Author contributions

**Sebastian Overmans**: Conceptualization, Microbial cultivations, Flow Cytometry, GC-MS analysis, Data curation, Data analysis, Visualization, Writing – original draft **Adair Gallo Jr**: Conceptualization, Particle preparation and characterization, Writing – review & editing

**Himanshu Mishra**: Conceptualization, Funding acquisition, Project administration, Resources, Methodology, Investigation, Supervision, Writing – review & editing

**Kyle J. Lauersen**: Conceptualization, Funding acquisition, Project administration, Resources, Methodology, Investigation, Supervision, Writing – review & editing

## Acknowledgments

The research reported in this publication was supported by KAUST baseline funding awarded to Kyle Lauersen and Himanshu Mishra. We would like to thank Yinfeng Xu and Peng Zhang from KAUST for assistance with the microparticle coating and environmental scanning electron microscopy, respectively. Figure 4 was created by Heno Hwang, scientific illustrator at King Abdullah University of Science and Technology (KAUST).

## References

[1] P.K. Ajikumar, K. Tyo, S. Carlsen, O. Mucha, T.H. Phon, G. Stephanopoulos, Terpenoids: opportunities for biosynthesis of natural product drugs using engineered microorganisms, Mol Pharm. 5(2) (2008) 167–90. 10.1021/mp700151b

[2] K.J. Lauersen, I. Huber, J. Wichmann, T. Baier, A. Leiter, V. Gaukel, V. Kartushin, A. Rattenholl, C. Steinweg, L. von Riesen, C. Posten, F. Gudermann, D. Lutkemeyer, J.H. Mussgnug, O. Kruse, Investigating the dynamics of recombinant protein secretion from a microalgal host, J Biotechnol. 215 (2015) 62–71. 10.1016/j.jbiotec.2015.05.001

[3] K.J. Lauersen, T. Baier, J. Wichmann, R. Wordenweber, J.H. Mussgnug, W. Hubner, T. Huser, O. Kruse, Efficient phototrophic production of a high-value sesquiterpenoid from the eukaryotic microalga *Chlamydomonas reinhardtii*, Metab Eng. 38 (2016) 331–343. 10.1016/j.ymben.2016.07.013

[4] E. Specht, S. Miyake-Stoner, S. Mayfield, Micro-algae come of age as a platform for recombinant protein production, Biotechnol Lett. 32(10) (2010) 1373–1383. 10.1007/s10529-010-0326-5

[5] T. Baier, J. Wichmann, O. Kruse, K.J. Lauersen, Intron-containing algal transgenes mediate efficient recombinant gene expression in the green microalga Chlamydomonas reinhardtii, Nucleic Acids Res. 46(13) (2018) 6909–6919. 10.1093/nar/gky532

[6] S. Gutiérrez, S. Overmans, G.B. Wellman, V.G. Samaras, C. Oviedo, M. Gede, G. Szekely, K.J. Lauersen, A synthetic biology and green bioprocess approach to recreate agarwood sesquiterpenoid mixtures, Green Chemistry. 26(5) (2024) 2577–2591. 10.1039/d3gc03708h

[7] M. Tashiro, H. Kiyota, S. Kawai-Noma, K. Saito, M. Ikeuchi, Y. Iijima, D. Umeno, Bacterial Production of Pinene by a Laboratory-Evolved Pinene-Synthase, Acs Synth Biol. 5(9) (2016) 1011–1020. 10.1021/acssynbio.6b00140

[8] F.K. Davies, V.H. Work, A.S. Beliaev, M.C. Posewitz, Engineering Limonene and Bisabolene Production in Wild Type and a Glycogen-Deficient Mutant of *Synechococcus* sp. PCC 7002, Front Bioeng Biotechnol. 2 (2014) 21. 10.3389/fbioe.2014.00021

[9] W.A. Duetz, H. Bouwmeester, J.B. van Beilen, B. Witholt, Biotransformation of limonene by bacteria, fungi, yeasts, and plants, Appl Microbiol Biotechnol. 61(4) (2003) 269–77. 10.1007/s00253-003-1221-y

[10] K.J. Lauersen, Eukaryotic microalgae as hosts for light-driven heterologous isoprenoid production, Planta. 249(1) (2019) 155–180. 10.1007/s00425-018-3048-x

[11] B.B. de Freitas, S. Overmans, J.S. Medina, P.Y. Hong, K.J. Lauersen, Biomass generation and heterologous isoprenoid milking from engineered microalgae grown in anaerobic membrane bioreactor effluent, Water Res. 229 (2023) 119486. 10.1016/j.watres.2022.119486

[12] S. Overmans, K.J. Lauersen, Biocompatible fluorocarbon liquid underlays for in situ extraction of isoprenoids from microbial cultures, Rsc Adv. 12(26) (2022) 16632–16639. 10.1039/d2ra01112c

[13] A.B. Lindstrom, M.J. Strynar, E.L. Libelo, Polyfluorinated Compounds: Past, Present, and Future, Environ Sci Technol. 45(19) (2011) 7954–7961. 10.1021/es2011622

[14] Y. Guan, Z. Liu, N. Yang, S. Yang, L.E. Quispe-Cardenas, J. Liu, Y. Yang, Near-complete destruction of PFAS in aqueous film-forming foam by integrated photo-electrochemical processes, Nature Water. 2(5) (2024) 443–452. 10.1038/s44221-024-00232-7

[15] B. Trang, Y. Li, X.S. Xue, M. Ateia, K.N. Houk, W.R. Dichtel, Low-temperature mineralization of perfluorocarboxylic acids, Science. 377(6608) (2022) 839-845. 10.1126/science.abm8868

[16] E. Poli, K.H. Jong, A. Hassanali, Charge transfer as a ubiquitous mechanism in determining the negative charge at hydrophobic interfaces, Nat Commun. 11(1) (2020) 901. 10.1038/s41467-020-14659-5

[17] B.R. Shrestha, S. Pillai, A. Santana, S.H. Donaldson, Jr., T.A. Pascal, H. Mishra, Nuclear Quantum Effects in Hydrophobic Nanoconfinement, J Phys Chem Lett. 10(18) (2019) 5530–5535. 10.1021/acs.jpclett.9b01835

[18] J. Nauruzbayeva, Z. Sun, A. Gallo, Jr., M. Ibrahim, J.C. Santamarina, H. Mishra, Electrification at water-hydrophobe interfaces, Nat Commun. 11(1) (2020) 5285. 10.1038/s41467-020-19054-8

[19] M.S. Sadullah, Y.F. Xu, S. Arunachalam, H. Mishra, Predicting droplet detachment force: Young-Dupré Model Fails, Young-Laplace Model Prevails, Communications Physics. 7(1) (2024) 89. 10.1038/s42005-024-01582-0

[20] J.N. Israelachvili, Intermolecular and surface forces, 3rd ed., Academic Press, Burlington, USA, 2010.

[21] P.-G. Gennes, F. Brochard-Wyart, D. Quéré, Capillarity and Wetting Phenomena: Drops, Bubbles, Pearls, Waves, Springer, New York, 2004.

[22] C.A. Quist-Jensen, F. Macedonio, C. Conidi, A. Cassano, S. Aljlil, O.A. Alharbi, E. Drioli, Direct contact membrane distillation for the concentration of clarified orange juice, Journal of Food Engineering. 187 (2016) 37–43. 10.1016/j.jfoodeng.2016.04.021

[23] A. Santana, A.S.F. Farinha, A.Z. Toraño, M. Ibrahim, H. Mishra, A first-principles approach for treating wastewaters, International Journal of Quantum Chemistry. 121(5) (2020) e26501. 10.1002/qua.26501

[24] H. Mishra, A. Farinha, S. Sinha, Functionalized SiO2 microspheres for extracting oil from produced water, Google Patents, 2018.

[25] A. Gallo, K. Odokonyero, M.A.A. Mousa, J. Reihmer, S. Al-Mashharawi, R. Marasco, E. Manalastas, M.J.L. Morton, D. Daffonchio, M.F. McCabe, M. Tester, H. Mishra, Nature-Inspired Superhydrophobic Sand Mulches Increase Agricultural Productivity and Water-Use Efficiency in Arid Regions, ACS Agricultural Science & Technology. 2(2) (2022) 276–288. 10.1021/acsagscitech.1c00148

[26] K. Odokonyero, A. Gallo, Jr., V. Dos Santos, H. Mishra, Effects of superhydrophobic sand mulching on evapotranspiration and phenotypic responses in tomato (*Solanum lycopersicum*) plants under normal and reduced irrigation, Plant Environ Interact. 3(2) (2022) 74–88. 10.1002/pei3.10074

[27] K. Odokonyero, A. Gallo, Jr., H. Mishra, Nature-inspired wax-coated jute bags for reducing post-harvest storage losses, Sci Rep. 11(1) (2021) 15354. 10.1038/s41598-021-93247-z

[28] J. Beekwilder, A. van Houwelingen, K. Cankar, A.D. van Dijk, R.M. de Jong, G. Stoopen, H. Bouwmeester, J. Achkar, T. Sonke, D. Bosch, Valencene synthase from the heartwood of Nootka cypress (*Callitropsis nootkatensis*) for biotechnological production of valencene, Plant Biotechnol J. 12(2) (2014) 174–82. 10.1111/pbi.12124

[29] A. Gallo, F. Tavares, R. Das, H. Mishra, How particle-particle and liquid-particle interactions govern the fate of evaporating liquid marbles, Soft Matter. 17(33) (2021) 7628–7644. 10.1039/d1sm00750e

[30] K.J. Lauersen, J. Wichmann, T. Baier, S.C. Kampranis, I. Pateraki, B.L. Moller, O. Kruse, Phototrophic production of heterologous diterpenoids and a hydroxy-functionalized derivative from *Chlamydomonas reinhardtii*, Metab Eng. 49 (2018) 116–127. 10.1016/j.ymben.2018.07.005

[31] M.N. Abdallah, G.B. Wellman, S. Overmans, K.J. Lauersen, Combinatorial Engineering Enables Photoautotrophic Growth in High Cell Density Phosphite-Buffered Media to Support Engineered *Chlamydomonas reinhardtii* Bio-Production Concepts, Front Microbiol. 13 (2022) 885840. 10.3389/fmicb.2022.885840

[32] D.S. Gorman, R. Levine, Cytochrome f and plastocyanin: their sequence in the photosynthetic electron transport chain of *Chlamydomonas reinhardi*, P Natl Acad Sci USA. 54(6) (1965) 1665

[33] M. Liu, Y.C. Lin, J.J. Guo, M.M. Du, X. Tao, B. Gao, M. Zhao, Y. Ma, F.Q. Wang, D.Z. Wei, High-Level Production of Sesquiterpene Patchoulol in *Saccharomyces cerevisiae*, Acs Synth Biol. 10(1) (2021) 158–172. 10.1021/acssynbio.0c00521

[34] Q.Q. Peng, Q. Guo, C. Chen, P. Song, Y.T. Wang, X.J. Ji, C. Ye, T.Q. Shi, High-Level Production of Patchoulol in *Yarrowia lipolytica* via Systematic Engineering Strategies, J Agric Food Chem. 71(11) (2023) 4638–4645. 10.1021/acs.jafc.3c00222

[35] L. Vargas-Estrada, S. Torres-Arellano, A. Longoria, D.M. Arias, P.U. Okoye, P.J. Sebastian, Role of nanoparticles on microalgal cultivation: A review, Fuel. 280 (2020) 118598. ARTN 118598 10.1016/j.fuel.2020.118598

[36] G. Cheloni, E. Marti, V.I. Slaveykova, Interactive effects of copper oxide nanoparticles and light to green alga *Chlamydomonas reinhardtii*, Aquat Toxicol. 170 (2016) 120–128. 10.1016/j.aquatox.2015.11.018

[37] C. Wang, C.Q. Lan, Effects of shear stress on microalgae - A review, Biotechnol Adv. 36(4) (2018) 986–1002. 10.1016/j.biotechadv.2018.03.001

[38] J. Neupert, D. Karcher, R. Bock, Generation of Chlamydomonas strains that efficiently express nuclear transgenes, Plant J. 57(6) (2009) 1140–50. 10.1111/j.1365-313X.2008.03746.x

[39] S. Overmans, G. Ignacz, A.K. Beke, J.J. Xu, P.E. Saikaly, G. Szekely, K.J. Lauersen, Continuous extraction and concentration of secreted metabolites from engineered microbes using membrane technology, Green Chemistry. 24(14) (2022) 5479–5489. 10.1039/d2gc00938b

[40] N. Subramanian, A. Qamar, A. Alsaadi, A. Gallo, Jr., M.G. Ridwan, J.G. Lee, S. Pillai, S. Arunachalam, D. Anjum, F. Sharipov, N. Ghaffour, H. Mishra, Evaluating the potential of superhydrophobic nanoporous alumina membranes for direct contact membrane distillation, J Colloid Interface Sci. 533 (2019) 723–732. 10.1016/j.jcis.2018.08.054

[41] Z.D. Hendren, J. Brant, M.R. Wiesner, Surface modification of nanostructured ceramic membranes for direct contact membrane distillation, J Membrane Sci. 331(1-2) (2009) 1–10. 10.1016/j.memsci.2008.11.038

[42] M. Rezaei, D.M. Warsinger, V.J. Lienhard, M.C. Duke, T. Matsuura, W.M. Samhaber, Wetting phenomena in membrane distillation: Mechanisms, reversal, and prevention, Water Res. 139 (2018) 329–352. 10.1016/j.watres.2018.03.058

[43] S. Pillai, A. Santana, R. Das, B.R. Shrestha, E. Manalastas, H. Mishra, A molecular to macro level assessment of direct contact membrane distillation for separating organics from water, J Membrane Sci. 608 (2020) 118140. 10.1016/j.memsci.2020.118140

[44] N. Seltenrich, From Drinking Water to Individual Body Burden: Modeling Toxicokinetics of Four PFAS, Environ Health Perspect. 131(1) (2023) 14001. 10.1289/EHP12514

